# Population density and temperature influence the return on maternal investment in wild house mice

**DOI:** 10.1101/2020.06.30.177089

**Authors:** Yannick Auclair, Nina Gerber, Barbara König, Anna K. Lindholm

**Affiliations:** Department of Evolutionary Biology and Environmental Studies, University of Zurich, Winterthurerstrasse 190, 8057 Zurich, Switzerland

**Keywords:** Body mass, Longevity, Maternal effects, Population density, Reproductive success, Weaning weight

## Abstract

In mammals, reproduction is influenced by competitive stress, temperature and food availability and these factors might be crucial already during early life. Favourable early life environment and high maternal investment are expected to improve survival and reproduction. In mammals, maternal investment via lactation predicts offspring growth. As body mass is often associated with fitness consequences, females have the potential to influence offspring fitness through their level of investment, which might interact with effects of population density and temperature. Here, we investigate the relationship between pup body mass at day 13 (used as approximation for weaning mass) and individual reproductive parameters as well as longevity under natural variation in population density and temperatures. Further, we assess the extent to which mothers influence the body mass of their offspring until weaning. We analysed life data of 384 house mice (*Mus musculus domesticus*) from a free-living wild population that was not food limited. We found a complex effect of population density, temperature and maternal investment on life-history traits related to fitness: Shorter longevity with increasing pup body mass at day 13; delayed reproduction of heavier pups when raised at warmer temperatures; and increased lifetime reproductive success for heavier pups at high densities. House mice could use population density and temperature as cues for predicting future environmental conditions, allowing a mother to adjust her investment according to the environment in which offspring will breed in order to maximise fitness. This study highlights the importance of considering ecological conditions in combination with maternal effects.

## Introduction

Reproductive success determines the fitness of an individual. In mammals, three factors universally influence reproduction: social stress, temperature and food availability (Bronson 1989). These factors might be not only crucial once maturity is reached, but also during early life. Early life is a critical period for newborns as any stress can have long-term irreversible consequences on their morphology (Lummaa and Clutton-Brock 2002; Tschirren et al. 2009), physiology (Mirescu et al. 2004; Sebaai et al. 2004), immunology (Edwards and Cooper 2005; Prager et al. 2010) or behaviour (Laviola and Terranova 1998; Levitsky and Barnes 1972; Lovic et al. 2001). Thus, conditions that are experienced during early life might shape future life-history strategies (Lindström 1999; Metcalfe and Monaghan 2001).

In mammals, females typically provide most if not all of the parental care offspring require to reach independence. Of the different aspects of care, lactation represents the most essential parental investment component that determines food availability for offspring (Gittleman 1985). Previous laboratory studies have shown that maternal investment through lactation can explain up to 65% of the variation observed in body mass at weaning in the house mouse *Mus domesticus* (Atchley and Zhu 1997; Cox et al. 1959; El Oksh et al. 1967). This maternal source of variance in offspring phenotype can be partitioned into prenatal maternal effects like resources allocated to an egg, and postnatal maternal effects such as maternal behaviour. In species where newborns have a prolonged period of maternal dependence until weaning, the contribution of postnatal maternal effects on offspring body size can outweigh that of prenatal maternal effects (Reinhold 2002; Steiger 2013). Since offspring body mass at weaning correlates with maternal investment (Seals: Don Bowen et al. 2001; Mice: Falconer 1947; Ferrari et al. 2015; McDowell et al. 1930; Review: Mateo 2009), weaning represents the best time point to assess cumulative maternal energy allocation (pre- and postnatal). Body mass at or near weaning represents a good alternative to direct metabolic measurements of parental investment that are often too complicated or invasive to be used on wild populations (König et al. 1988; Sadowska et al. 2013).

There are many examples of benefits of being relatively heavier from small mammals to large herbivores. Heavier individuals usually have a higher probability of settlement in their population (Wauters et al. 1993), can achieve higher dominance ranks (e.g Klemme et al. 2006; Krackow 1993), reproduce earlier and/or produce more offspring (e.g Anderson and Fedak 1985; Dobson and Michener 1995; Festa-Bianchet et al. 2000; Fuchs 1982), or survive better (e.g Millar and Hickling 1990; Murie and Boag 1984; Wauters and Dhondt 1989). Moreover, body mass is considered a reliable proxy for the quality or health of an individual (Oli and Dobson 2003; Peters 1986). The influence of size on performance is not only observed during adulthood but can also be detected at earlier life stages (Dias and Marshall 2010). The positive relationship between offspring and adult body mass (Birgersson and Ekvall 1997; Festa-Bianchet et al. 2000) and the increased offspring quality at weaning observed whenever offspring received extended maternal care (Dahle and Swenson 2003) suggest that higher maternal investment improves offspring fitness.

However, there are situations in which investment in bigger offspring is not favourable, as under certain conditions smaller offspring size might have benefits or because investing in bigger offspring is costly for mothers (Dantzer 2013). Mothers can regulate their investment during lactation through milk quality and may invest differently in offspring depending on litter size, condition or offspring sex. In house mice for example, investment into individual offspring (and their weaning weight) decreases with increasing litter size (König et al. 1988). Different maternal investment depending on the sex has been demonstrated in mammals as sexes benefit differently from maternal investment (Caecero et al. 2018 for calves; Landete-Castillejos et al. 2005 for red deer; Quesnel et al. 2017 for kangaroos). Likewise, other offspring traits or condition can influence the return on maternal investment. For example, the timing of birth can influence growth, survival, and fitness (Clutton-Brock et al. 1982) and thus the conditions at birth might influence the returns of maternal investment. Population density is strongly related to social stress (Clutton-Brock 1988; Gaillard et al. 1997; Gilbert and Krebs 1991; Saitoh 1981) and the population density and/or climatic conditions (temperature) during gestation and at birth of a litter might influence physiological needs of the mother as well as the offspring, and affect future reproductive behaviour. Modification of maternal care according to the social and seasonal conditions experienced when offspring are born might allow the adaptive adjustment of offspring mass to the conditions expected once sexually mature (Dantzer et al. 2013).

Here, we aim to investigate the influence of social cues (population density) and the physical environment (temperature) in combination with maternal investment on reproductive success using long-term data from an intensively monitored wild population of house mice (*Mus musculus domesticus*) in middle Europe that is not limited by food. This reflects a natural situation for house mice in Europe and North America, as they normally breed in man-made structures with non-limited food sources (Bronson 1979). House mice are nonhibernating and are not strictly seasonal breeders.

They produce several litters over an extended breeding period that may cover even up to 12 months per year (König and Lindholm 2012), as they can breed at freezing temperatures as long as enough food is available (Bronson and Pryor 1983). First, we test whether pup body mass at day 13 (which we used as an approximation for weaning weight) has an effect on the age at first reproduction, longevity and lifetime reproductive success (LRS), while accounting for population density and temperature at birth as well as sex differences. We predict increasing LRS and longevity with pup body mass. Further, we expect that at unfavourable conditions (low temperature/high population density) reproduction is reduced and that under these conditions, first reproduction is delayed. Second, we assess the maternal influence on pup body mass while controlling for offspring sex as well as litter size and litter sex ratio. Last, we assessed the relationship between the estimated weaning mass and adult body mass. This study offers the opportunity to analyse the flexibility and the evolutionary consequences of maternal care dependent on environmental conditions, given a natural annual cycle in temperature and fluctuation in local population density.

## Materials and methods

### Study population

Data were collected from a free-living house mouse population in a 70m^2^agricultural building in Illnau, Switzerland. Although mice could easily exit the barn through numerous gaps, none of the large mammalian and avian predators that occur outside could enter. This reflects a normal situation for house mice, since they typically breed out of reach of nest predators (Latham and Mason 2004) and large predators as cats, dogs or foxes are usually not efficient enough to control a population (Timm 1994). The high permeability of the building towards mice did not allow us to directly monitor exits and entrances (but see Runge and Lindholm 2018 for estimation of migration propensity). Water and food, a 50/50 mixture of oats and hamster food (Landi AG, Switzerland), were provided *ad libitum* to match conditions under which natural house mouse populations are typically observed in Western Europe (Berry 1970). The entire population inside the building was captured on average every seven weeks to estimate adult population density and to examine animals.

### Reproductive activity

Reproduction occurred in 40 artificial nest boxes. We searched for new litters approximately every ten days between January 2007 and December 2009. Each new litter was given an identification number. Pups were sexed according to their anogenital distance and genital morphology (Hotchkiss and Vandenbergh 2005) and aged according to morphological development (but not by body mass). Skin pigmentation, development of the ears, the growth of the fur, teeth eruption, and eye development give reliable cues about the age of the pups (Figure S1, the day of birth was considered as day 1). Besides these regular 10-day checks, we also checked nest boxes whenever pups raised inside were forecast to be 13 days of age. Experimenters gained experience with sex and age estimates in the laboratory where births of litters were precisely documented. Because birth in house mice can last several hours, an uncertainty of about one day remains in our age estimation.

### Body mass measurements

Pups were weighed to the nearest 0.1g when they were 13 days old (±1 day). This age is the last day before weaning when they can be reliably captured as they are still blind, largely unreactive and entirely dependent on milk (actual weaning starts at day 17 and is completed at approximately 21-23 days old (König 1993; König and Markl 1987)). We would have preferred to obtain body mass at the onset of weaning, but pups could only be safely caught up to 13 days of age as they become mobile and may run away to evade capture from age 14 days on (own observations; see also Mikesic and Drickamer 1992). Pup body mass is expected to increase linearly between day 13 and day 17 (Bronson 1979; König and Markl 1987). This increase, however, is likely to stop between day 17 and day 21 as females encourage their pups to eat solid food until they are independent (König and Markl 1987). Using data collected in a laboratory study using descendants from our house mouse population (Ferrari et al. 2015), we found that body mass at day 13 was a strong predictor of body mass at the onset of weaning at day 17 (*r*_207_ = 0.85, p < 0.001; Figure S2). We, therefore, consider mass at day 13 as a “weaning mass estimate”.

In captures of the entire population, each individual was weighed to the nearest 0.1g, and those weighing at least 17-18g were considered adults (Pelikán 1981, Auclair et al. 2014). At that age, adults were individually marked with RFID tags (Trovan® ID-100A implantable micro transponder: 0.1 g weight, 11.5 mm length, 2.1 mm diameter; implanter Trovan® IID100E; Euro ID Identifikationssysteme GmbH and Co, Germany) for other research projects (e.g. Auclair et al. 2014; Harrison et al. 2018; Ferrari et al. 2019). Because the population is captured every seven weeks and some resident animals may have been outside of the building, individuals differ in the age at which they were first captured as adults.

### Genotyping and parentage analysis

An ear tissue sample was collected from every pup that was weighed at 13 days of age, every adult that was tagged, and on all corpses found. Following the same procedure as in Auclair et al. (2014), DNA was amplified using 25 microsatellite loci and a parentage analysis allowed assignment of the mother and the father of each individual to a 95% level of confidence using Cervus 3.0 (Marshall et al. 1998). Only fully assigned offspring and corpses that gave good quality DNA were kept in the analysis. Litter sizes were estimated by counting the number of pups with the same estimated day of birth (± 1 day) that were assigned to the same mother. This was necessary as litters often share nest boxes, and are sometimes relocated to other nest boxes (Ferrari et al. 2019).

### Life-history traits

Individual reproductive success was assessed using genetic parentage analyses and separately defined as both age at first reproduction, and the total number of offspring weaned (defined as surviving to 13 days of age) throughout life. The total number of offspring weaned was monitored for 384 pups that did not emigrate as juveniles (167 females and 217 males) born between January 2007 and December 2009 from 178 litters produced by 120 mothers. Among these 384 individuals, 219 (100 females and 119 males) reproduced, so their age at first reproduction was available. They produced a total of 2631 pups. The collection of corpses also allowed us to calculate longevity for 147 (56 females and 91 males) of these 384 individuals. We included LRS and life expectancy measures of these mice until February 2012, the last time a living focal mouse was recorded. The adult population density was estimated using an algorithm developed by Runge and Lindholm (2018) that allowed us to calculate the density of adults at any given date. The temperature was measured by a thermologger (HOBO U12-013) installed on an inside wall and mean monthly temperatures were used for the analyses.

### Statistical analyses

Statistical tests were carried out using R 3.5.1 (R Development Core Team 2013). We estimated the influence of mother identity, population density, litter size, sex, and sex ratio on pup body mass using a linear mixed-effects model (Bates et al. 2012). Mother identity was defined as a random factor while population density, sex ratio, litter size, sex and their interactions were defined as fixed factors. Final models were selected by dropping insignificant two-way interactions from the full model (model including fixed effects with all 2-way interactions) as long as the model fit did not decrease. We proceeded with this approach until the AIC of the model no longer decreased (an ΔAIC of 2 was considered a model improvement) or no non-significant interactions were left. The influence of mother identity on the adult body mass of the offspring was assessed with a separate linear mixed-effects model with mother identity defined as a random factor and population density, pup body mass, sex, litter size and their interaction defined as fixed factors. Continuous predictors were centred and model assumptions were checked visually and they were met for all models. We also did not detect any under- or overdispersion of the GLMM. The importance of mother identity was estimated as the proportion of the total variance explained by the random factor “mother identity” and tested by a likelihood ratio test comparing this model to a generalised least square model having the same fixed factors structure but no random factor (Zuur et al. 2009). To investigate significant interaction effects with continuous predictors, we checked whether the 95% confidence intervals of the slopes predicted by the model overlapped (Aiken and West 1991). For the illustration and slope analysis of continuous by continuous variable interactions, one of the predictors was categorised with the 0.5 quantile for sex ratio or the 0.25 and 0.75 quantile for population density, weaning weight and temperature.

The influence of pup body mass at day 13 on longevity was analysed with a linear model accounting for pup body mass, sex, population density, temperature and their interactions. The number of offspring weaned was analysed with a zero-inflated model with the same fixed effects structure (Zeileis et al. 2008). For the age at first reproduction, we performed a Cox proportional hazard regression (Kaplan and Meier 1958), again with the same structure for fixed effects. This analysis considers that the timing of reproduction varies among groups, and whether the event of reproduction happens or not. For all models, the model assumptions were tested visually.

## Results

### Life-history consequences of pup body mass at day 13

The age at first reproduction was influenced by the temperature during the month of birth and its interaction with pup body mass at day 13 (Table 1, Figure 1). When raised at cold temperatures, mice reproduced earlier in their life compared to offspring born at warmer temperatures. Temperature was, however, strongly associated with population density, with population density being generally higher in summer when mean monthly temperatures were higher (ρ = 0.676, Figure 2). Despite the significant interaction between the temperature during the month of birth and pup body mass, the effect of pup body mass was not significantly different from zero, neither at high nor low temperatures considering that the 95% confidence intervals are overlapping (Figure 1).

**Table 1.** Influence of pup body mass at day 13, sex, population density and temperature at birth and their interactions on individuals’ life-history traits (measured by the age at first reproduction), lifetime reproductive success (analysing total number of offspring raised until day 13), the probability to reproduce at all, using the binomial & logit part of zeroinflated regression model and longevity. VIF_PopDens_=2.168; VIF_Temp_=2.160.

| Life-history traits<br>(Response) | Fixed Effects | $\beta$ | SE | t-value |
| --- | --- | --- | --- | --- |
| Age at first reproduction<br>CoxPH<br>N = 384<br>(number of events 219) | **Temperature at birth** | **0.322** | **0.139** | **2.310** |
|  | Pup body mass | -0.058 | 0.250 | -0.231 |
|  | Sex (Male) | 0.315 | 0.334 | 0.944 |
|  | Population density | -0.006 | 0.010 | -0.618 |
|  | **Temperature at birth : Pup body mass** | **-0.043** | **0.016** | **-2.648** |
|  | Temperature at birth : Sex(Male) | -0.029 | 0.029 | -0.998 |
|  | Temperature at birth : Population density | -0.001 | 0.000 | -1.833 |
|  | Pup body mass : Population density | 0.002 | 0.001 | 1.432 |
| Lifetime reproductive<br>success<br>(# offspring weaned)<br>logit<br>Zero Inflated Regression<br>(neg binomial)<br>N = 384 | **Intercept** | **27.716** | **11.133** | **2.490** |
|  | **Pup body mass** | **-3.926** | **1.561** | **-2.516** |
|  | **Population density** | **-5.129** | **2.169** | **-2.364** |
|  | Temperature at birth | 0.501 | 0.347 | 1.442 |
|  | Sex (Male) | -0.009 | 0.149 | -0.058 |
|  | **Pup body mass : Population density** | **0.794** | **0.311** | **2.552** |
|  | Pup body mass : Temperature at birth | -0.022 | 0.017 | -1.314 |
|  | Population density : Temperature at birth | -0.065 | 0.056 | -1.152 |
| Lifetime reproductive<br>success<br>(Reproducing or not)<br>binomial<br>Zero Inflated Regression<br>(neg binomial)<br>N = 384 | Intercept | -1.382 | 21.751 | -0.064 |
|  | Pup body mass | -0.119 | 3.061 | -0.039 |
|  | Population density | 0.222 | 4.302 | 0.052 |
|  | Temperature at birth | 0.378 | 0.651 | 0.581 |
|  | Sex (Male) | 0.106 | 0.261 | 0.407 |
|  | Pup body mass : Population density | -0.020 | 0.617 | -0.032 |
|  | Pup body mass : Temperature at birth | 0.028 | 0.039 | 0.708 |
|  | Population density : Temperature at birth | -0.087 | 0.105 | -0.833 |
| Longevity<br>N = 147 | **Intercept** | **1299.000** | **425.400** | **3.054** |
|  | **Pup body mass** | **-123.800** | **55.040** | **-2.249** |
|  | **Sex (male)** | **-488.400** | **225.300** | **-2.167** |
|  | Population density | -2.963 | 2.353 | -1.259 |
|  | Temperature at birth | 20.960 | 27.050 | 0.775 |
|  | Pup body mass : Sex | 35.070 | 30.820 | 1.138 |
|  | Pup body mass : Population density | 0.464 | 0.296 | 1.566 |
|  | Pup body mass : Temperature at birth | -3.261 | 3.172 | -1.028 |
|  | Sex : Temperature at birth | 0.167 | 0.698 | 0.238 |
|  | Sex (male) : Population density | 3.793 | 7.405 | 0.512 |
|  | Population density : Temperature at birth | -0.006 | 0.050 | -0.123 |

**Figure 1.**
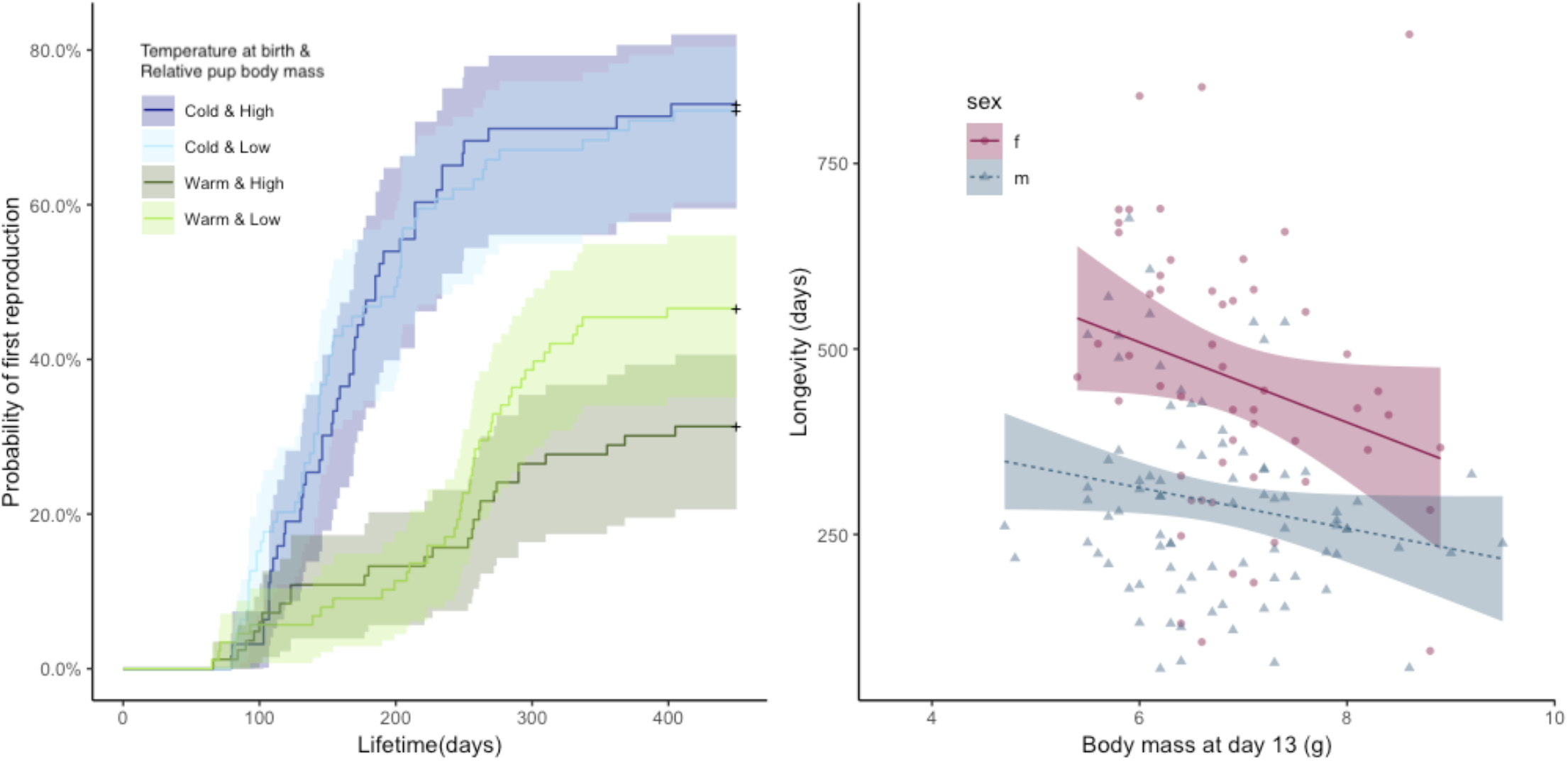
Probability of reproduction (left) and longevity (right) with pup body mass at day 13 and relevant interaction. The shaded areas indicate the 95% confidence intervals. Temperature at birth and pup body mass at day 13 were categorised for illustration. Values above the median were considered as high, respectively warm and values below the median as low, respectively cold.

**Figure 2.**
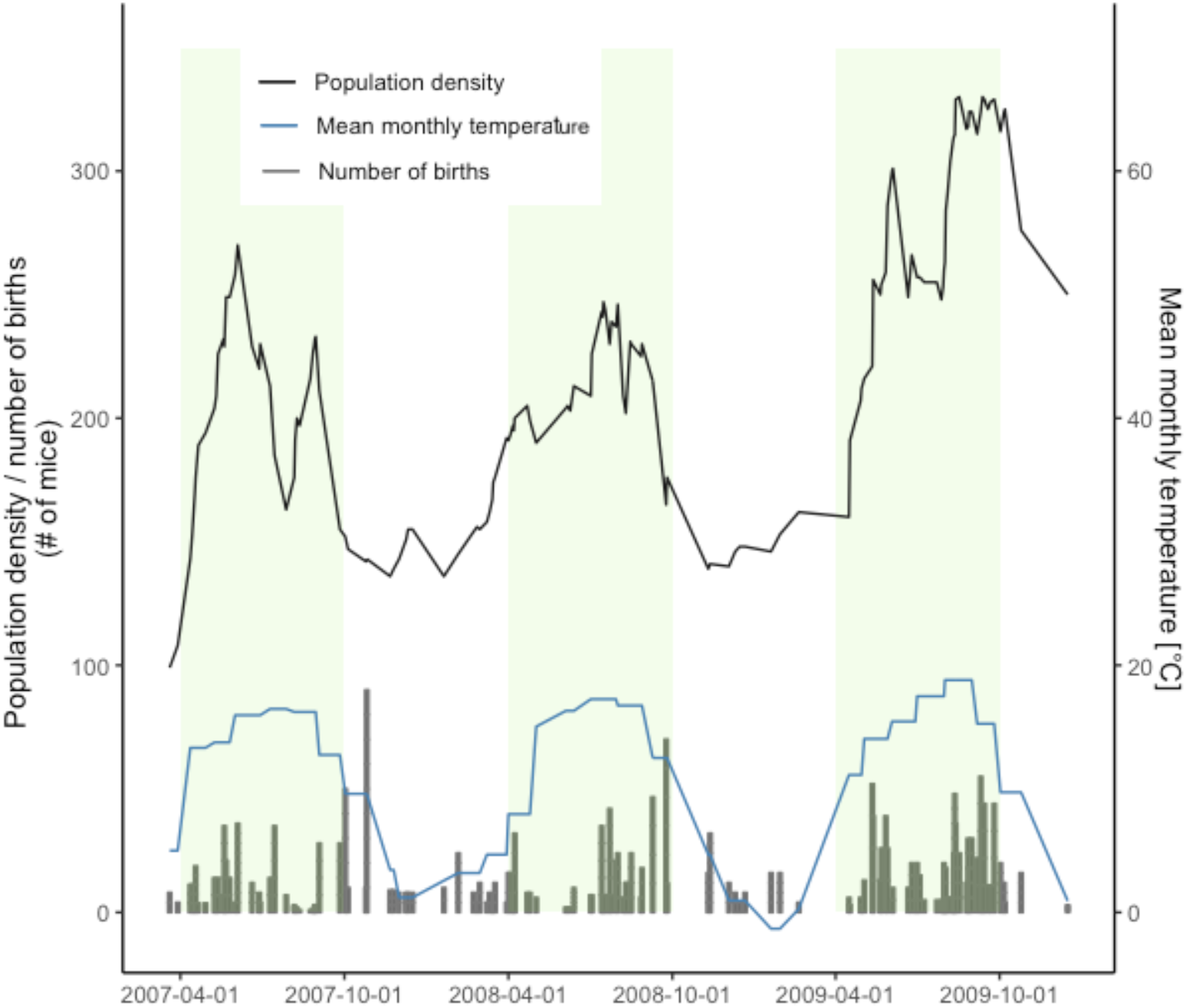
Population density, mean monthly temperature in the barn and number of births over time. Green shaded areas represent the summer months and white areas referring to winter months.

Individual lifetime reproductive success (LRS), as measured by the number of offspring surviving until day 13 over a lifetime, was influenced by the interaction of pup body mass at day 13 with population density at birth (Figure 3, Table 2). There was a positive effect of pup body mass on LRS at high population densities, but the slopes did not differ significantly from zero at low densities (Figure 3). As 165/384 individuals did not breed at all, we also tested for factors predicting breeding. When looking at the binomial part of the zero-inflated GLM, it seems that the effect of pup body mass and density on LRS stem from differences in the number of offspring, rather than differences in the probability of reproducing (Table 2).

**Figure 3.**
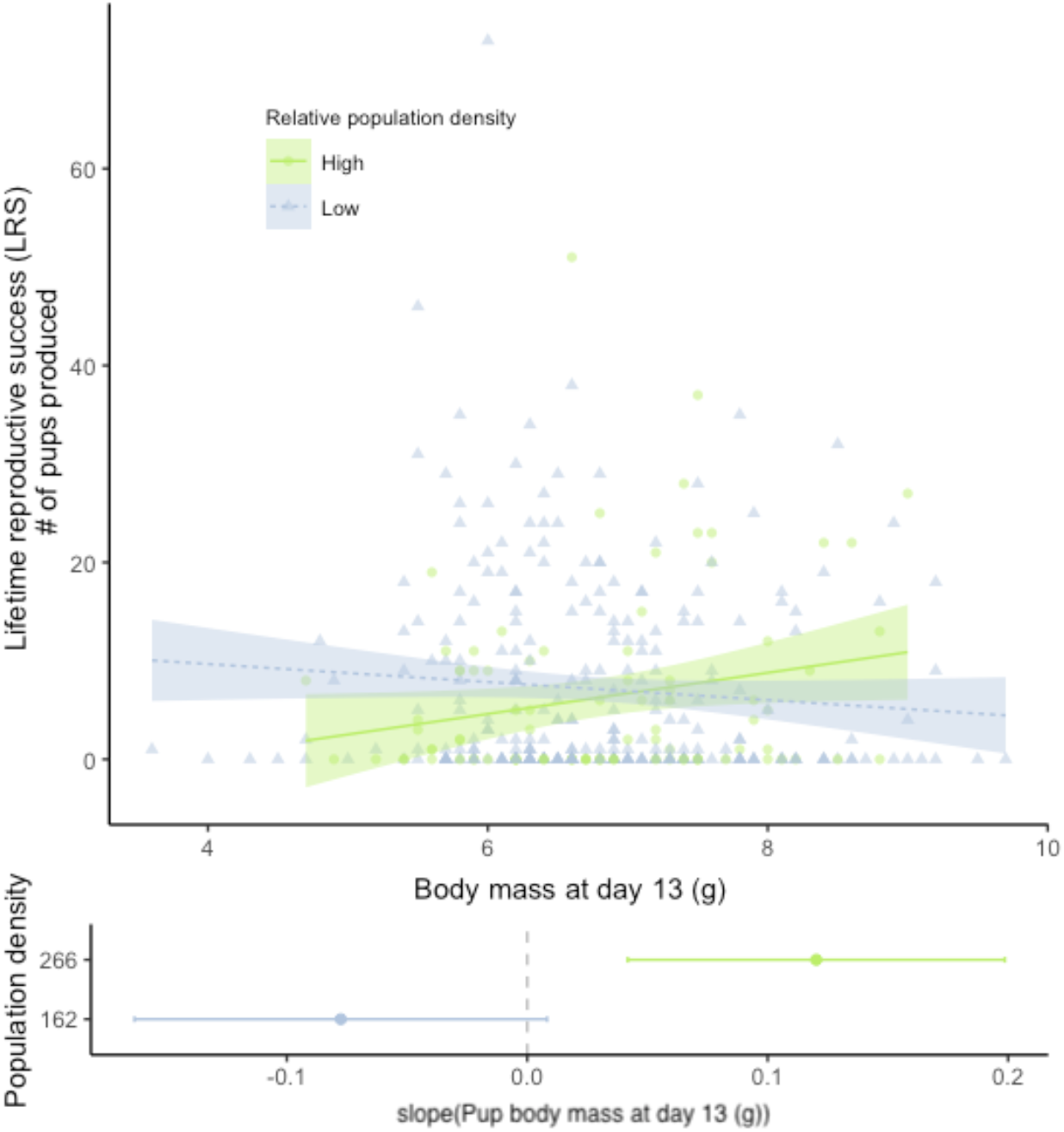
Relationship of lifetime reproductive success (LRS) with pup body mass at day 13. The shaded areas indicate the 95% confidence intervals. Population density was categorised for the figure and the slope analysis with high population density above the 0.75 quantile and low population density below the 0.25 quantile.

Longevity was influenced by sex, with females living longer than males (Table 3; Figure 1), and pup mass at day 13. Pup body mass had a significant negative influence on longevity (Table 3; Figure 1). Neither population density nor the time in the year nor any interactions had a significant effect on longevity (Table 3). There was no bias in the sex ratio of the 237 individuals for which we found no dead bodies (and thus we were not able to analyse their longevity) and assumed they died outside of the building (111 females vs 126 males; *χ^2^*= 0.95, df = 1, *p* = 0.330).

### Influence of mother identity, sex, sex ratio, litter size and population density on pup body mass

Differences between mothers accounted for 48% of the total variance observed in pup body mass at day 13 when corrected for litter size (Table 4). When not corrected for litter size, the mother’s contribution accounted for 49% of the variance in pup body mass (Table S1). Pup body mass was significantly influenced by the interaction of sex with population density (Table 4). Female pup body mass was positively related to population density, whereas the slope for male body mass did not differ from zero, considering 95% confidence intervals (Figure 4). Pup body mass at day 13 was positively related to the temperature during the month a litter was born (Figure 4, Table 4). Pup body mass was also influenced by the interaction between litter size and sex ratio (Table 4), such that slopes of male- and female-biased sex ratios differed from each other, but the slopes did not differ from zero at male- or female-biased sex ratio (Figure 4), thus, this effect may not be biologically relevant.

**Table 2.** Influence of mother identity, sex, sex ratio, and litter size on pup body mass at day 13 (*N* = 368); estimated by linear mixed model with “ML”. Model coefficients of the main effects are reported for the model excluding interaction terms.

| Pup body mass |  |  |  |  |
| --- | --- | --- | --- | --- |
| Fixed effects: | B | SE | t-value | P |
| (Intercept) | 6.646 | 0.367 | 18.096 | <0.001 |
| Litter size | -0.099 | 0.049 | -2.021 | 0.044 |
| Sex (male) | 0.776 | 0.312 | 2.490 | 0.014 |
| Sex ratio | -0.902 | 0.321 | -2.805 | 0.006 |
| Population density | 0.001 | 0.002 | 0.533 | 0.594 |
| Temperature at birth | 0.026 | 0.013 | 2.009 | 0.046 |
| Litter size : sex ratio | 0.189 | 0.067 | 2.810 | 0.005 |
| Sex (male) : Population density | -0.003 | 0.001 | -2.320 | 0.021 |
| $\omega^2 = 0.585$ | | | | |
| $R^2 = 0.617$ | | | | |
| Random effects: | SD | Residual |  |  |
| Mother identity | 0.671 | 0.698 |  |  |

**Figure 4.**
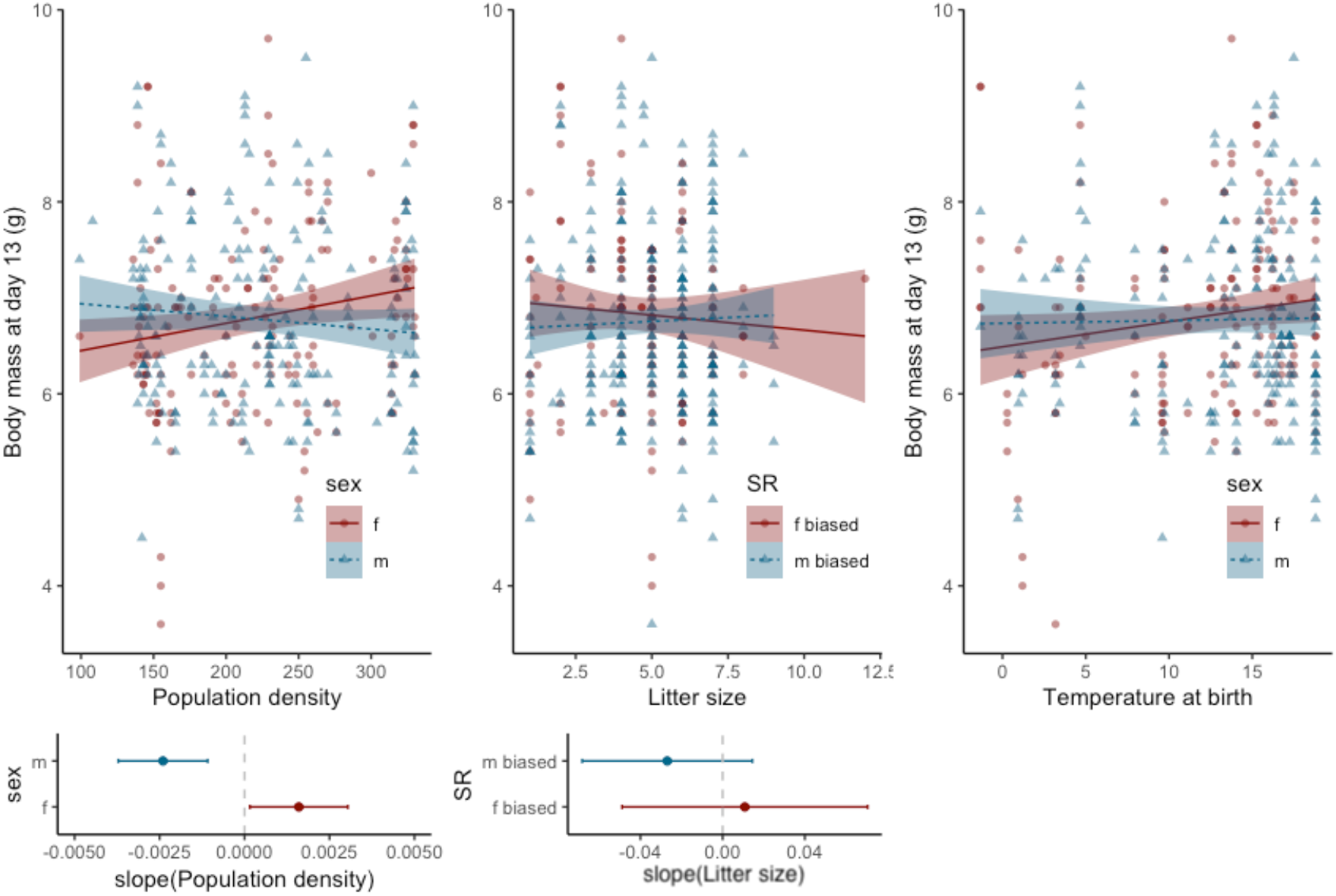
Relationship of population density (left), litter size (middle) and temperature (right) with pup body mass at day 13; the shaded area indicates the 95% confidence intervals. Sex ratio (SR) was categorized and was considered male biased when greater than 0.5 and female biased when below 0.5. Population density was categorised for the figure and the slope analysis as high whenever above the 0.75 quantile and as low population density below the 0.25 quantile. Figures below show slope analysis with 95% confidence intervals.

### Influences of mother identity, pup body mass, and sex on adult body mass

As adult body mass increased with age, the measure of body mass had to be corrected for the age when adults were first weighed, which varied (see Figure S3). A negative exponential function was fitted to the data for both sexes, and the residuals were used as an index for adult body mass in further analyses.

Once adult, the influence of mother identity decreased to 12% of the total variance observed in the body mass when first captured as adults (Table S2). The effect of pup body mass at day 13 on adult body mass depended on the population density at birth, as well as the sex of the offspring (Figure S4, Table S2). Thus, at high population density, pup body mass in females was positively related to adult body mass, whereas there was no effect at low population density. In males, pup body mass was related to adult mass regardless of density (Figure S4). There was a significant interaction of the litter size with pup body mass. However, the slopes at different weaning weights were not different from zero (Figure S5). We have also found a significant interaction between the temperature at birth and the offspring sex, but again, simple slope analysis revealed that neither of the slopes differed from zero.

## Discussion

This study aimed to examine the lifetime fitness consequences of variation in offspring body mass at day 13 (used as an approximation for weaning mass) under varying social and climatic conditions during the month of birth, and to investigate the extent to which lactating mothers influence the body mass of their offspring. Here, we report delayed reproduction when pups were raised at warmer temperatures, shorter longevity with increasing pup body mass at day 13, and a positive effect of body mass on lifetime reproductive success for pups born at high population densities. Furthermore, we observed that mothers had a significant influence on offspring body mass accounting for nearly half of all residual variation.

### Effect of pup body mass, population density and temperature on life-history traits

In our study population longevity decreased with increasing pup body mass at day 13. We had predicted increased survival with body mass, as it is known to be correlated with body fat, which increases survival during harsh periods (e.g. McMahon et al. 2015, Ronget et al. 2018). However, in our system, heavier pups died younger, in both males and females. The shorter lifespan of pups with a higher body mass might be due to strong intra-sexual competition acting in both sexes (Miller et al. 2002). Heavy offspring may experience more frequent agonistic interactions that may compromise their survival compared to their smaller counterparts. They may be perceived as a threat and attacked by dominant individuals and their higher likelihood to become dominant increases their risk to be injured or killed while being repeatedly challenged by same-sex conspecifics (Oakeshott 1974). Social competition may, however, differ between sexes (Stockley et al. 2013). For instance, the evolution of physiological suppression in females may help to decrease the rate of intrasexual aggression (Drickamer 1977; Kruczek et al. 1989) and may contribute to their survival advantage over males (Clutton-Brock 2009; Clutton-Brock et al. 1979; Clutton-Brock and Isvaran 2007). Alternatively, low mass offspring at high density may instead benefit from higher longevity, that allows them to survive until the next low density period, coinciding with a warm breeding period in spring. This is supported by the interaction between body mass and temperature on the probability of reproduction (Table 1) and the negative relationship between mass and longevity (Figure 1).

We expected that a heavier weaning mass would allow an earlier onset of reproduction hence leading to a higher reproductive success if individuals manage to reproduce regularly (Roff 2002). The effect of pup body mass at day 13 on age at first reproduction was dependent on the mean monthly temperature at birth, but neither at high nor at low temperature was this difference significantly different from zero. However, individuals born at high temperatures delayed reproduction. Summer (when temperatures were high) was the main breeding season and was when population density was highest (see Figure S5). The effect of temperature in our model might have masked an effect of population density. Thus, the delayed reproduction when raised during warmer periods could have been due to an increase in reproductive competition or a reduction in the number of available nesting sites (Gilbert and Krebs 1991; Jorgenson et al. 1993; Komers et al. 1997; Saitoh 1981). Additionally, high population density is associated with stress, which inhibits sexual maturity (Van Zegeren 1980, Kruczek et al. 1989) and can cause reproduction to cease (Yasukawa et al. 1985). Previous studies have, however, indicated that low temperatures are associated with delayed puberty and reduced reproduction (Biggers et al. 1958, Barnett 1965) and breeding in mice is greatly reduced in winter (Bronson 2009, König and Lindholm 2012, Runge and Lindholm 2018). Individuals who are poor competitors are thus more likely to defer their reproduction and queue to acquire breeding or social positions (Kokko and Johnstone 1999; van de Pol et al. 2007). For those born in winter, the delay to the next warmer period that also has low density (spring) is short.

Pup body mass at day 13 only affected lifetime reproductive success (LRS) when the offspring were born at high population densities. Being born at high population densities might indicate deteriorating conditions. In our study population, high density at birth could predict harsher periods at sexual maturity, where heavier individuals have higher chances to breed. In many taxa, larger individuals have higher reproductive success (Dias and Marshall 2010). In mammals, bigger males typically have higher chances to acquire a breeding territory, become dominant and reproduce (Anderson and Fedak 1985; Bouteiller-Reuter and Perrin 2005; Clutton-Brock et al. 1979; Oakeshott 1974). Such advantages towards big individuals are not limited to males as intra-sexual competition also occurs within females (Clutton-Brock 2016; König and Lindholm 2012; Stockley and Bro-Jørgensen 2011). Female competition may occur to control territories housing the best nest sites or to control access to food (Bujalska 1973; Ostfeld 1985; Reimer and Petras 1967; Wolff 1993). Similarly to males, reproductive success can be skewed toward dominant females (Clutton-Brock et al. 1984; Rusu and Krackow 2004). Laboratory studies of mice have shown that a higher weaning weight in females results in larger litters and shorter inter-birth intervals during their first two reproductive events (Fuchs 1982).

A potential bias in our study is that we only had data for the reproductive success of non-dispersing individuals, which could influence the interpretation of the results if body mass influences the propensity to disperse (see Massot et al. 2002, Bonte and De La Peña 2009). In our study population, few individuals migrated among groups within the building, and pup body mass at age 13 days did not predict emigration from the building (Runge and Lindholm 2018). We, therefore, do not expect a bias related to a missing size class. Furthermore, house mice mainly disperse as subadults, before the onset of reproduction (Gerlach 1996), and our study only includes mice that were recorded as adults. Thus, we assume that individuals in our data set were non-dispersers.

### Early life population density and sex-specific effects of maternal care

Our results suggest that mothers adapt their investment in offspring according to the social environment. The social environment has been shown to influence offspring fitness (e.g. Siracusa et al. 2017). In our population, the social environment fluctuated within an individual’s life, and thus mothers may have enhanced the fitness of their offspring by adjusting offspring phenotypes to match the environment they will experience in the future. For such maternal effects to be adaptive, mothers need to be able to predict relevant aspects of the future environments of their offspring (Dantzer et al., 2013; Schwabl et al. 1997). When population growth is density dependent, population density can act as an indicator of conditions and reproductive output of the offspring. Thus, population density may act as a cue for adaptive reproductive adjustments in anticipation of density-dependent natural selection on offspring phenotypes (Dantzer et al. 2013).

At high population density, adult body mass was positively related to pup body mass at day 13 for males and females. At low population density, however, we did not find an effect of pup body mass on adult body mass in females. Thus, any reproductive benefits gained from higher adult body mass were only related to pup body mass at high population densities. Females seem to be able to compensate for low pup body mass at low population densities, where low competition for resources might increase the potential for compensatory growth (Sundström et al. 2013). Density-dependent maternal effects have been reported mainly in avian systems, where mothers adapt androgen allocation to the current social density (Remeš et al. 2011, Mazuc et al. 2003). Recently, similar effects of population density have been recorded in crickets, where females adjusted hormonal levels of their eggs to the experienced social environment (Crocker and Hunter 2018). Also, in mammals, population density was shown to alter maternal investment and increase offspring growth at high population densities to increase the probability of surviving winter periods. At low population density, on the other hand, fast growth is not favoured. This leads to conditions where the evolution of plastic maternal investment is favoured, whereby increased offspring growth coincides with high population density under which it enhances fitness (Dantzer et al. 2013).

Our results suggest that mothers increased their investment in heavier offspring at high densities only for female pups, but not for males. Similarly, female great tits increase androgen concentrations at high social densities (Remeš et al. 2011), which increase male, but not female growth rate. In mammals, milk production is energetically expensive, increasing females’ basal metabolic rate by up to 7.2 times (Hammond and Diamond 1992; König et al. 1988). Reducing investment in milk, leading to smaller offspring, could thus benefit mothers, if their fitness benefits outweigh fitness losses due to lighter offspring (as lower LRS of light offspring when born at low densities). Reproductive success was independent of pup body mass at day 13 at low densities, but at high population densities, pup body mass was positively related to lifetime reproductive success. The relationship of pup body mass to adult body mass differed between males and females. Unlike in males, this difference was positive at high densities for females. Sex differences could potentially be due to differences in maternal care, or differences in offspring feeding behaviour. Sex-specific resource allocation in house mice is unlikely because the immobile arched posture of nursing females limits their control of offspring access to the nipples (König 1989a). However, mothers might not increase investment to single offspring, but they might increase their investment in female-biased litters at high population densities. Furthermore, male and female pups may differ in their behaviour or metabolism (Garel et al. 2009) as seen in spotted hyenas *Crocuta crocuta* (Golla et al. 1999) and sea lions *Zalophus californianus* (Ono and Boness 1996), leading to different growth potential under competition. An inter-sibling competition favouring the access to milk in one sex or a sex differential digesting efficiency has, however, never been reported in house mice.

The ready access to food in the study population might have substantially lowered the costs of maternal investment. However, our study reflects natural conditions for house mice in Western Europe, as house mice usually live commensally with humans. Populations grow wherever food is easily accessible and available in good quantity, feral populations, that leave independent of humans all year round, are restricted to islands (Berry 1970; Latham and Mason 2004; Pocock et al. 2004). Thus, the food availability in our study reflects normal conditions, and we generally expect commensal populations to experience similar resource conditions as our study population.

## Conclusions

Our results show a strong effect of population density, temperature and maternal investment on life-history traits related to fitness. Similar to our results, other studies also find a more complex effect of pup body mass on fitness measures than often predicted (Ylönen et al. 2004, Pigeon and Pelletier 2018). We show that the effects of maternal investment are not independent of the effects of the environment. Our data provide evidence that house mice could use population density and temperature as cues for predicting future environmental conditions. This would allow a mother to adjust her investment according to the environment in which offspring will breed in order to maximise their fitness. Future experiments with controlled conditions along with the analysis of data from different years will help to unravel the complexity we find.

## Acknowledgments

We thank Gabi Stichel who transpondered the mice, Jari Garbely who performed the genetic analyses, and Corinne Ackermann for the parentage analysis and everyone who contributed to the data collection. We also thank Pawel Kotej and Marco Festa-Bianchet for helpful comments on the manuscript.

## Data accessibility

The data will be stored on Dryad upon acceptance of the Manuscript.

