## Supplementary material for "Population density and temperature influence the return on maternal investment in wild house mice": SupportingInformation_MaternalInvestmentOecologia.pdf

**Figure S1** Morphological development of a wild house mouse pup between day 1 and day 12 (picture courtesy of Manuela Ferrari) used for age assessment. Day 12 and day 13 (not pictured) differ in ear and hair development.

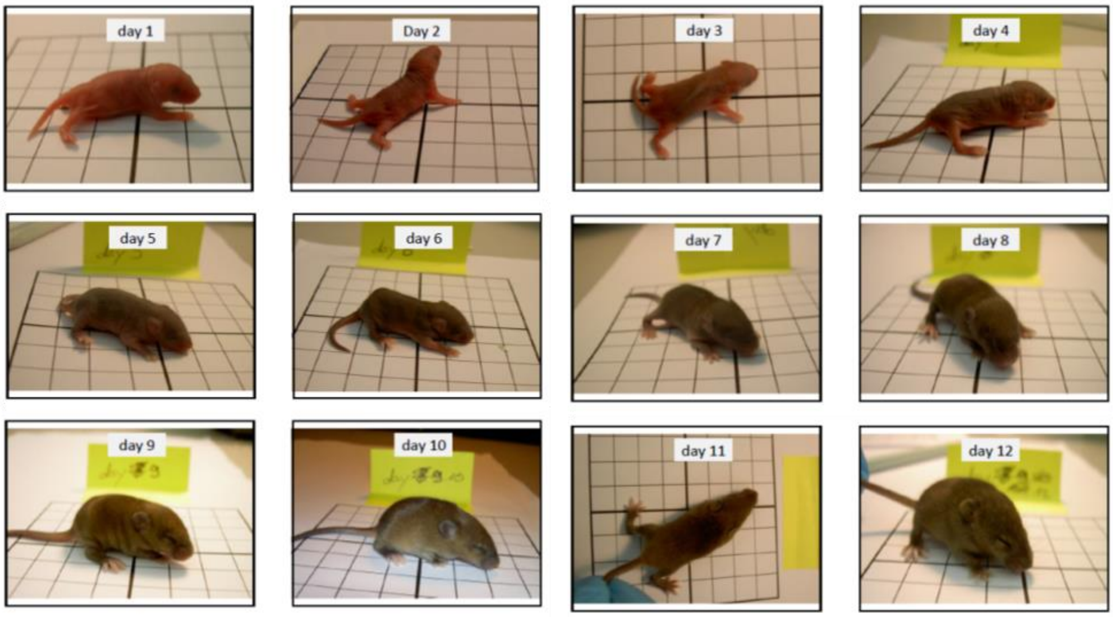

**Figure S2** Relationship of weight at day 13 and weight at the onset of weaning at day 17 (data from Ferrari et al. 2015).

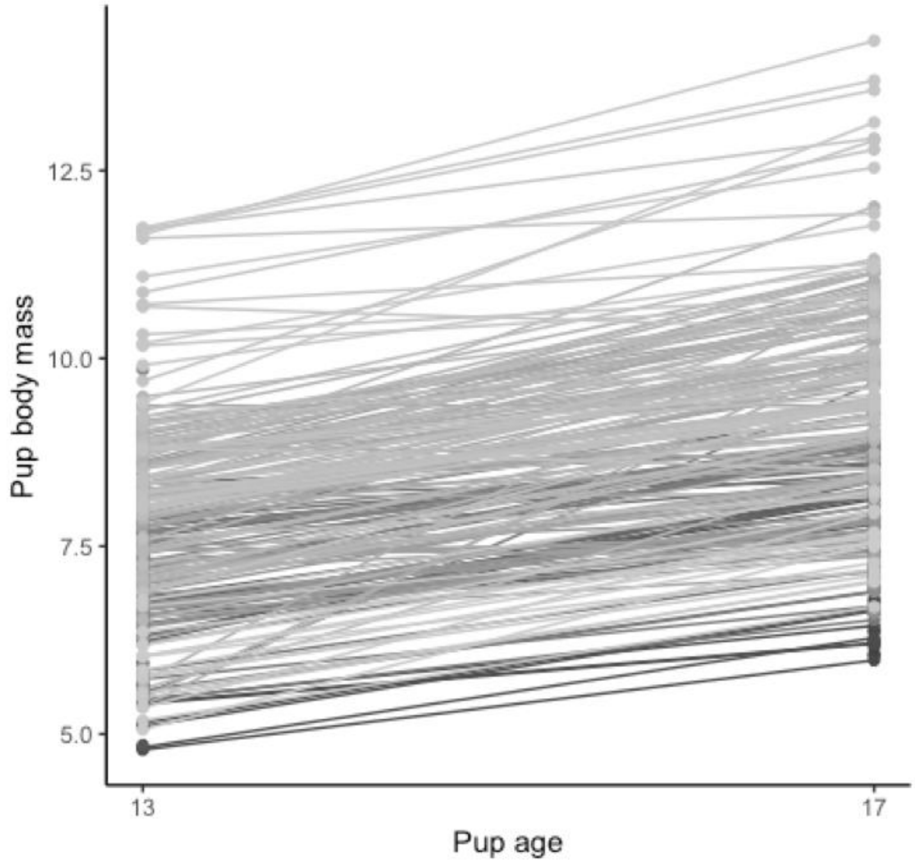

**Figure S3** Relationship of the age when mice were captured and their weight. The blue line shows the fitted negative exponential regression. The residuals have been used as “adult body mass” for further analyses.

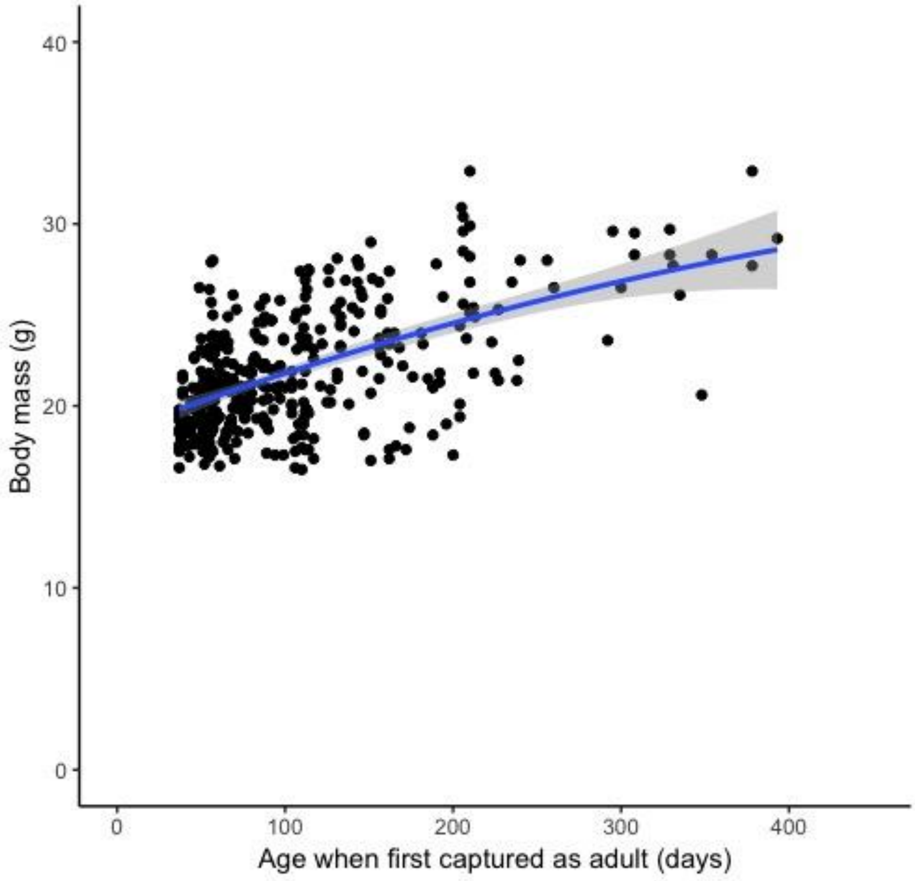

**Figure S4** Relationship of pup body mass and residuals of adult body mass (from a regression by age, see Figure S3). The shaded areas indicate the 95% confidence intervals. Figures below show slope analysis with 95% confidence intervals. Population density was categorised for the figure and the slope analysis as high whenever above the 0.75 quantile and as low whenever below the 0.25 quantile.

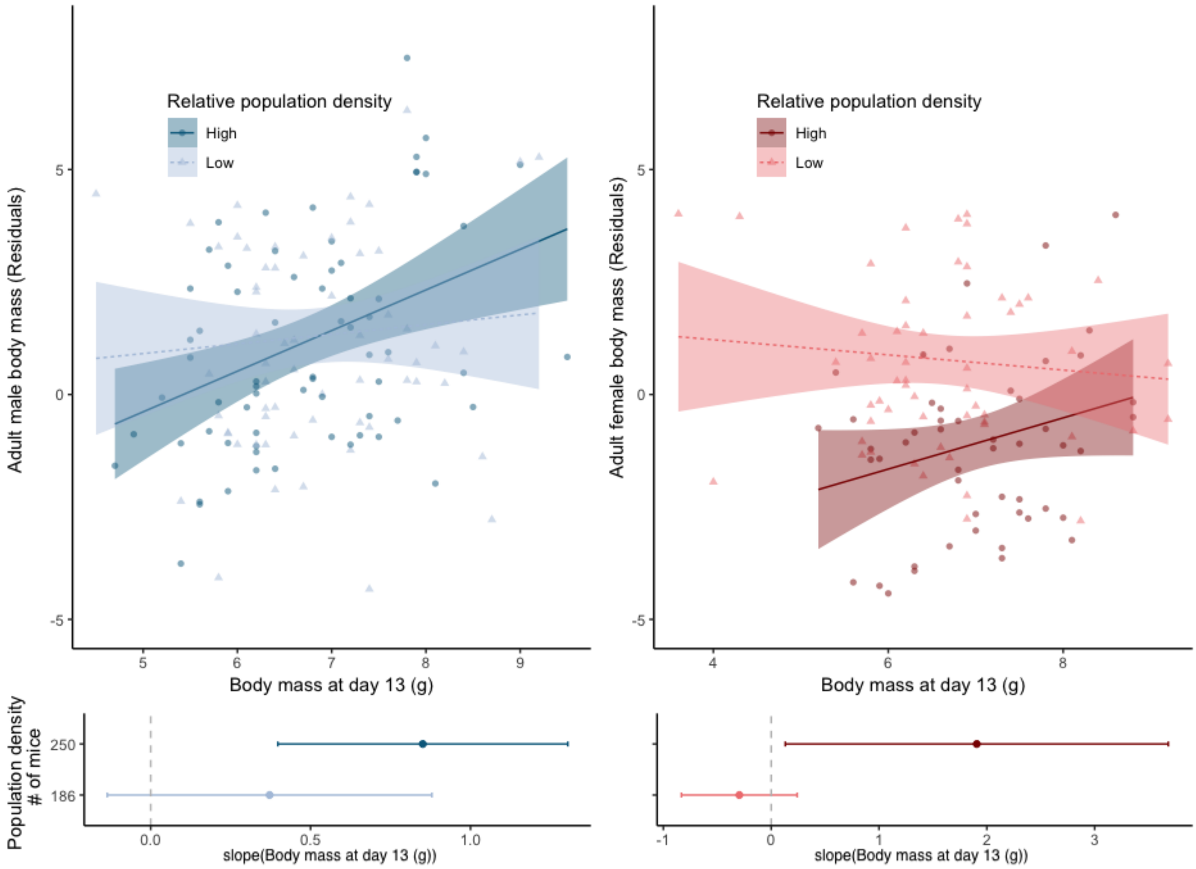

**Figure S5** Interaction between litter size and pup body mass. Pup body mass was categorised for the figure and the slope analysis as high (H) whenever above the 0.75 quantile and as low (L) whenever below the 0.25 quantile.

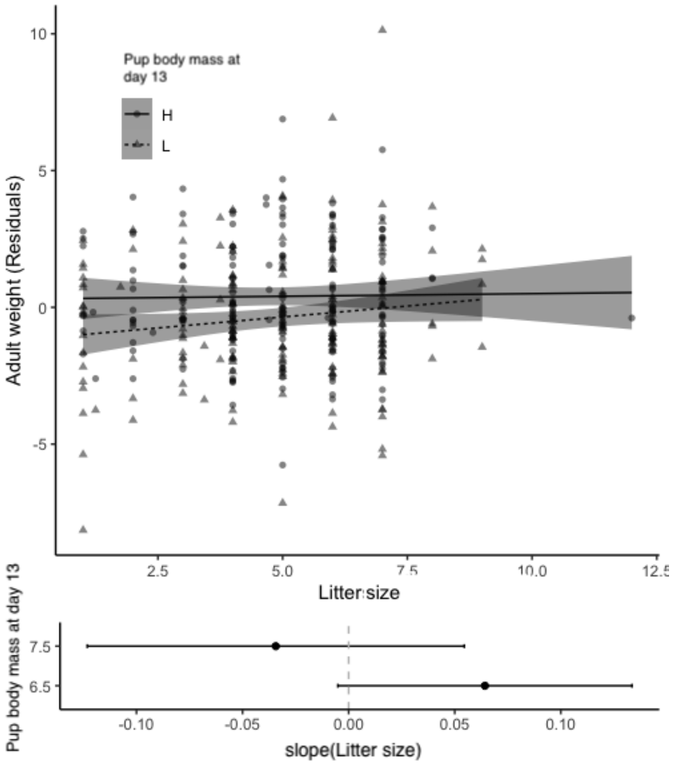

**Table S1.** Effects of influence of mother identity, sex, without the effect of litter size on pup body mass ($N$ = 368); estimated by linear mixed model with "ML". The model coefficient of the main effects are reported for the model excluding interaction terms.

| **Pup body mass** | | | | |
|---|---|---|---|---|
| Fixed effects: | ß | SE | t-value | p |
| **(Intercept)** | **5.842** | **0.379** | **15.429** | **<0.001** |
| Sex (male) | 0.374 | 0.256 | 1.464 | 0.145 |
| **Population density** | **0.005** | **0.002** | **2.265** | **0.024** |
| Temperature at birth | -0.008 | 0.020 | -0.398 | 0.691 |
| **Sex (male): Population density** | **-0.011** | **0.003** | **-3.736** | **<0.001** |
| $\omega^2 = 0.585$ | | | | |
| $R^2 = 0.634$ | | | | |
| Random effects: | SD | Residual | | |
| Mother identity | 0.703 | 0.695 | | |

**Table S2.** Effects of mother identity, pup body mass (pup bm), sex and their interactions on adult body mass (*N* = 368); estimated by linear mixed model with maximum likelihood estimation.

| **Adult body mass** | | | | |
|---|---|---|---|---|
| Fixed effects: | ß | SE | t-value | p |
| **(Intercept)** | **-6.874** | **2.312** | **-2.974** | **0.003** |
| **Pup body mass** | **1.225** | **0.352** | **3.479** | **0.001** |
| Sex(males) | 0.576 | 0.824 | 0.698 | 0.486 |
| Sex Ratio | -0.778 | 0.498 | -1.563 | 0.120 |
| **Litter size** | **1.188** | **0.471** | **2.523** | **0.012** |
| Temperature at birth | 0.020 | 0.043 | 0.457 | 0.648 |
| **Pup body mass : Litter size** | **-0.181** | **0.071** | **-2.563** | **0.011** |
| **Sex (Males) : Temperature at birth** | **-0.149** | **0.055** | **-2.704** | **0.007** |
| **Pup body mass : Sex(females) : Population density** | **-0.002** | **0.001** | **-3.522** | **0.001** |
| **Pup body mass: Sex(males) : Population density** | **0.001** | **0.001** | **2.009** | **0.046** |

$\omega^2 = 0.326$

$R^2 = 0.405$

| Random effects: | SD | Residual |
|---|---|---|
| Mother identity | 0.771 | 2.118 |
